# Substrate stress relaxation regulates monolayer fluidity and leader cell formation for collectively migrating epithelia

**DOI:** 10.1101/2024.08.26.609529

**Authors:** Frank Charbonier, Junqin Zhu, Raleigh Slyman, Cole Allan, Ovijit Chaudhuri

**Author notes:** **Corresponding author:** Ovijit Chaudhuri, **Email:**. **Author Contributions:** F.C. and O.C designed research; F.C., J.Z., and R.S. performed research, F.C, J.Z., C.A., and R.S. contributed new reagents/analytic tools, F.C., J.Z., and C.A. analyzed data, and F.C. and O.C. wrote the paper. **Competing Interest Statement:** The authors declare no competing interests.

## Abstract

Collective migration of epithelial tissues is a critical feature of developmental morphogenesis and tissue homeostasis. Coherent motion of cell collectives requires large scale coordination of motion and force generation and is influenced by mechanical properties of the underlying substrate. While tissue viscoelasticity is a ubiquitous feature of biological tissues, its role in mediating collective cell migration is unclear. Here, we have investigated the impact of substrate stress relaxation on the migration of micropatterned epithelial monolayers. Epithelial monolayers exhibit faster collective migration on viscoelastic alginate substrates with slower relaxation timescales, which are more elastic, relative to substrates with faster stress relaxation, which exhibit more viscous loss. Faster migration on slow-relaxing substrates is associated with reduced substrate deformation, greater monolayer fluidity, and enhanced leader cell formation. In contrast, monolayers on fast-relaxing substrates generate substantial substrate deformations and are more jammed within the bulk, with reduced formation of transient lamellipodial protrusions past the monolayer edge leading to slower overall expansion. This work reveals features of collective epithelial dynamics on soft, viscoelastic materials and adds to our understanding of cell-substrate interactions at the tissue scale.

**Significance Statement:** Groups of cells must coordinate their movements in order to sculpt organs during development and maintain tissues. The mechanical properties of the underlying substrate on which cells reside are known to influence key aspects of single and collective cell migration. Despite being a nearly universal feature of biological tissues, the role of viscoelasticity (i.e., fluid-like and solid-like behavior) in collective cell migration is unclear. Using tunable engineered biomaterials, we demonstrate that sheets of epithelial cells display enhanced migration on slower-relaxing (more elastic) substrates relative to faster-relaxing (more viscous) substrates. Building our understanding of tissue-substrate interactions and collective cell dynamics provides insights into approaches for tissue engineering and regenerative medicine, and therapeutic interventions to promote health and treat disease.

## Introduction

Collective cell migration is a fundamental process in development, tissue homeostasis, wound healing, and cancer metastasis. (1, 2) Tissue repair and morphogenesis require coordinated motion and force generation between cells for efficient and cohesive movement of epithelial sheets. (3, 4) Such cooperation is enabled by mechanical coupling of cells to each other through cell-cell adhesions and to the underlying substrate through cell-matrix adhesions. Collective interactions between individual cells can lead to emergent phenomena not observed in single cells, such as long-ranged force transmission and coherent flows within tissues. (5–9) The leading edge of freely expanding epithelial monolayers is typically characterized by large, highly motile cells (i.e., leader cells) at the front of finger-like protrusions, as well as contractile actomyosin cables spanning multiple cell lengths. (10–13) Spatiotemporal coordination of these distinct migration modes (i.e., lamellipodium-mediated cell crawling and purse-string contraction of cables) allows monolayers to migrate across heterogenous substrates and efficiently close wounds. (14–17)

Given the long-standing recognition that physical properties of the extracellular matrix (ECM) regulate cellular behaviors ranging from single cell spreading and motility to proliferation and differentiation, increasing focus has been directed towards the role of cell-substrate interactions and tissue mechanics in collective migration. (18–22) In particular, migration of epithelial monolayers in 2D is generally faster on stiffer substrates and can even be directed by gradients of stiffness (i.e., collective durotaxis). (23, 24) In-vitro studies on collective epithelial dynamics have typically used rigid (i.e., plastic, glass) or soft elastic substrates. Linearly elastic materials behave like a spring, deforming instantaneously in proportion to an applied load and returning to their original shape after removal of the load. A cell applying traction forces to a purely elastic material will experience resistance to deformation that is constant in time. In contrast, most biological tissues and ECMs are viscoelastic. (25, 26) Viscoelastic materials exhibit time-dependent mechanical properties and dissipate energy like viscous liquids. A cell applying traction forces to a viscoelastic material will experience decreasing resistance to deformation, due to a behavior known as stress relaxation, as energy is dissipated through matrix rearrangement and flow or other mechanisms. Substrate viscoelasticity is known to regulate cell adhesion, force generation, stem cell differentiation, proliferation, apoptosis, morphogenetic processes, matrix deposition, and single cell migration. (27–31) For example, several studies indicate that cells spread to a greater extent when cultured on more viscoelastic substrates that exhibit faster stress relaxation. (32, 33) In the context of migration of single cells, adherent cells exhibit faster migration with enhanced stress relaxation, with filopodia-mediated migration of single cells on the faster relaxing substrates. (34) However, it is unclear how these single cell phenomena translate to the behavior of multicellular collectives on viscoelastic substrates.

Here we use viscoelastic biomaterials to investigate the role of substrate stress relaxation in 2D collective cell migration. We pattern MDCK monolayers on engineered alginate hydrogels with independently tunable stiffness and stress relaxation, then analyze cell motion, coordination, and cytoskeletal organization to study collective behaviors. We find that epithelial clusters on slower-relaxing substrates are more motile and expand faster than clusters on faster-relaxing substrates. These differences are attributed to increased leader cell formation, increased monolayer fluidity, and decreased substrate deformation.

## Results

### Patterning epithelial monolayers on tunable viscoelastic substrates

To address the role of substrate viscoelasticity in collective cell migration, we fabricated thin alginate hydrogels with tunable stress relaxation. While unmodified alginate polymers are bioinert, cell-adhesive ligands such as the RGD (Arg-Gly-Asp) peptide sequence can be covalently coupled to the alginate chain to promote integrin-based cell adhesion. (35, 36) Alginate hydrogels are formed by ionic crosslinking with divalent cations such as Ca^2+^. Decreasing the length (i.e., molecular weight) of alginate polymer results in hydrogels with faster stress relaxation, while increasing the alginate molecular weight yields slower-relaxing hydrogels. Ionic crosslinking density can be modulated to maintain the same initial elastic modulus or stiffness. Further, diffusion studies indicate hydrogels formed in such a manner have similar pore sizes. (37) Thus, stress relaxation in alginate substrates can be tuned independently of the initial elastic modulus (i.e., stiffness), ligand density, and pore size. (38) This alginate hydrogel system has been used to study cell adhesion and spreading in 2D, as well as cell spreading, proliferation, migration, and differentiation in 3D. (27, 30, 32, 39–41) We generate alginate hydrogels with an initial elastic modulus (stiffness) of ∼20kPa and an order of magnitude difference in relaxation halftime (∼45s for fast-relaxing vs ∼700s for slow-relaxing). (**Fig. 1 A-C)** The stiffness of these hydrogels is similar to that of elastic substrates used in recent 2D studies of collective epithelial migration, while the stress relaxation overlaps with the biologically relevant range and is on timescales relevant to processes involved in cell motility (e.g., adhesion formation and extension of lamellipodial protrusions). (23, 25, 27, 34, 42) We next adapted a method for generating micropatterned epithelial clusters on 2D substrates to study collective migration of expanding monolayers. In this model wound healing assay, a PDMS stencil placed on the cell culture substrate is used to confine confluent monolayers to defined regions.(23, 43, 44) Upon removal of the stencil, the tissues expand outwards into the free space. In contrast to glass, plastic, and soft linearly elastic materials (e.g., polyacrylamide) used in similar studies, our alginate substrates exhibit viscoelastic creep under static loading. Therefore, careful optimization of the stencil properties and cell seeding procedures was required in order to generate successful clusters without mechanical disruption of the hydrogel substrate. (see Methods and Materials). Using this approach, we generated rectangular monolayers (500 μm wide) or circular monolayers (750 μm diameter) on thin alginate-RGD hydrogels and tracked tissue expansion over 24+ hours with live confocal timelapse imaging. **(Fig. 1D)**

**Figure 1.**
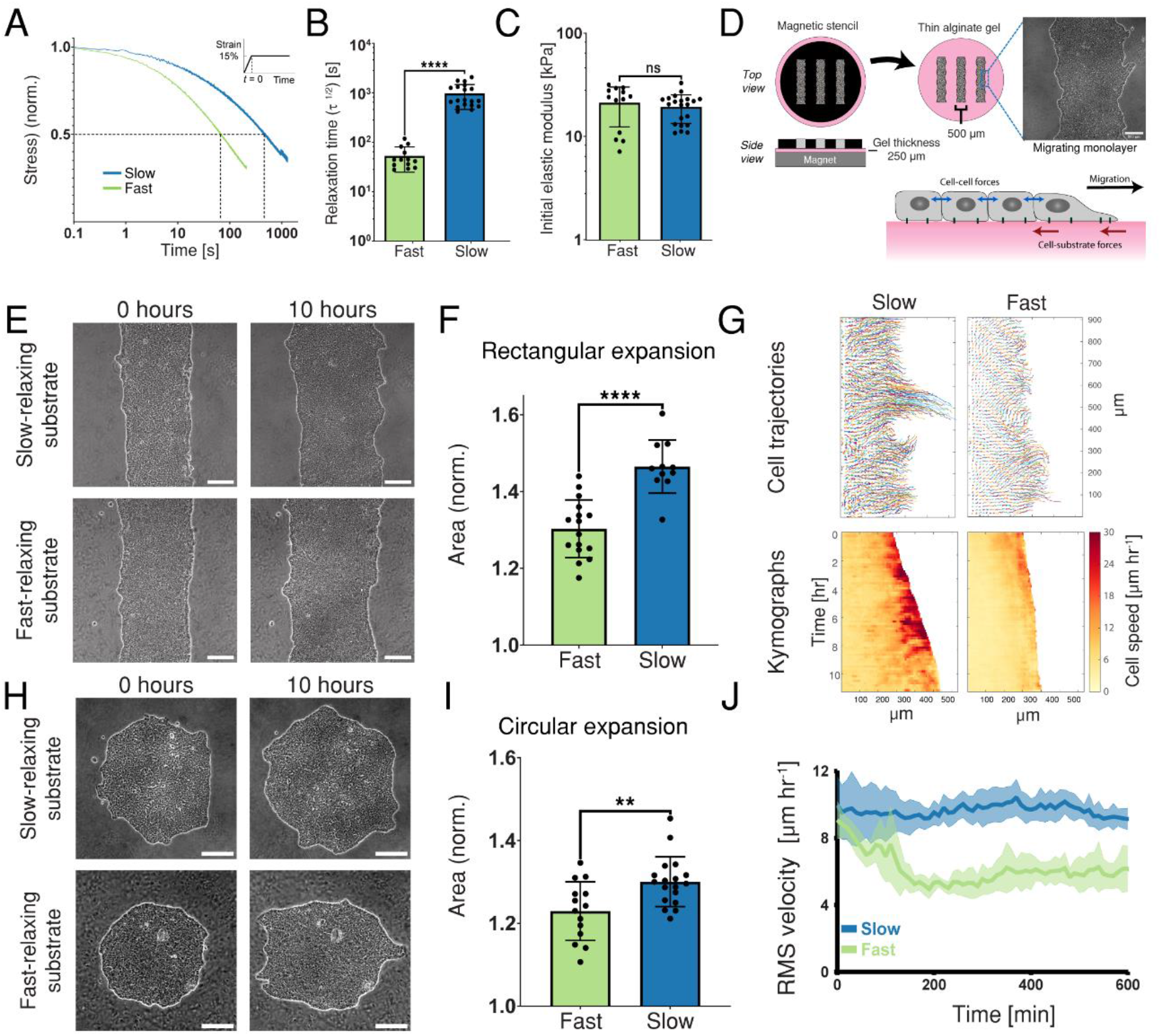
Epithelial monolayers expand faster on slow-relaxing vs fast-relaxing substrates. **A)** Representative stress relaxation profiles for fast-relaxing and slow-relaxing hydrogels prepared from alginate polymers with different molecular weights. **B)** Relaxation timescale values for fast-relaxing and slow-relaxing hydrogels. (n=13/19 for fast/slow, respectively). Two-tailed Mann-Whitney test, p<0.0001 **C)** Initial elastic modulus values from the same set of alginate gels as in B. (n=13/21 for fast/slow, respectively). Two-tailed Mann-Whitney test, p=0.3259. **D)** Schematic of micropatterning assay. Monolayers with defined geometry are generated on 2D alginate substrates and migration is tracked over 24+ hours. **E**,**H)** Sample images of rectangular **(E)** and circular **(H)** monolayers on fast- and slow-relaxing substrates at 0 hours and 10 hours after the start of imaging. **F**,**I)** Monolayer area 10hrs post-stencil-removal for rectangular (F) and circular (I) assay format, normalized by starting area. **F)** (n=16/12 for fast/slow, respectively, from two independent experiments). Two-tailed t-test with Welch’s correction, p<0.0001. **I)** (n=14/18 for fast/slow, respectively, from two independent experiments). Two-tailed t-test with Welch’s correction, p<0.0047. **G)** Cell trajectories and kymographs for rectangular clusters over 10+ hours of migration, from particle image velocimetry (see methods). **J)** Root mean square velocity over entire monolayer area for 10 hours of migration. Circular assay format. (n=7/10 clusters for fast/slow, respectively). Error bars represent SD for all panels.

### Epithelial monolayers expand faster on slow-relaxing vs fast-relaxing substrates

We find that MDCK monolayers exhibit more rapid migration on slow-relaxing substrates relative to fast-relaxing substrates with the same initial elastic modulus. For both rectangular clusters migrating outwards along a single axis **(Fig. 1 E and F)** and for circular clusters expanding radially **(Fig. 1 H and I)**, the expansion of area is faster on the slow-relaxing substrates. From particle image velocimetry (PIV) applied to phase contrast images we can obtain velocity fields over the entire monolayer. (7, 45) Regions of faster migration are largely restricted to the monolayer edge, especially for fast-relaxing substrates (**Fig. 1G)**. Further, while migration velocities are similar at the beginning of the migration assay, within an hour, clear differences emerge between the average migration speed of cells on slow relaxing versus fast relaxing substrates, with cells on slow-relaxing substrates exhibiting a higher root mean square average velocity (**Fig. 1J**). Together, these data indicate that epithelial monolayers exhibit faster collective migration on slower relaxing substrates.

### Leader cell formation is enhanced on slow-relaxing substrates

To investigate how these differences in collective migration arise, we first looked at the leading edge of the expanding monolayers. In line with prior studies, the leading edge does not move forward as a uniform front but roughens through formation of finger-like instabilities. (43, 46) Cells along the monolayer periphery periodically extend lamellipodia-like protrusions past the leading edge. (**Fig. 2A**). These cells will sometimes surge forward to become leaders at the front of multicellular fingers, whereas other times these protrusions will retract. Protrusive leader cells drive localized and transient bursts of faster migration (**Fig. 2B, Movie S1**), which are visible in kymographs of the leading edge as interspersed periods of steeper and shallower slope (**Fig. 2 C**). To examine whether the dynamics of leader cell formation shifted on substrates with different relaxation timescales, we tracked the emergence and duration of transient protrusions along monolayer edges over a 20-hour period of migration **(Movie S2)**. Protrusive leader cells form more frequently and migrate faster on slow-relaxing as compared to fast-relaxing hydrogels, whereas leader cell lifetime and persistence are similar in both cases (**Fig. 2 D-G**). On a per-cluster basis, greater numbers of leader cells correlate strongly with faster tissue expansion over 10 hours. (**Fig. 2H)**. Thus, these data imply that substrate viscoelasticity impacts collective migration by regulating leader cell formation.

**Figure 2.**
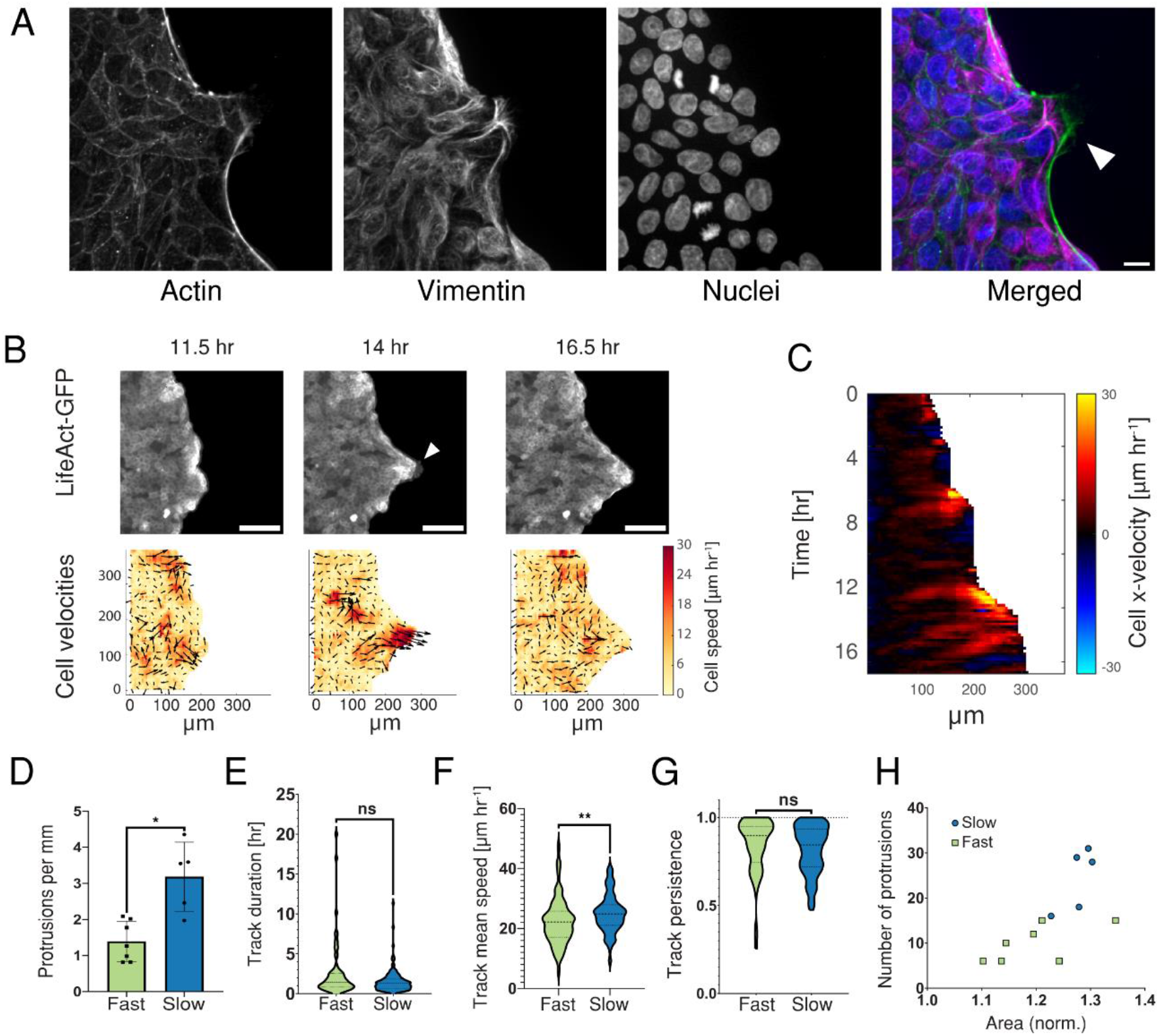
Enhanced leader cell activity on substrates with slow relaxation. **a)** Immunofluorescence image of leader cell (white arrow) forming at the monolayer edge. (Scale bar, 10µm.) **B)** Representative example of transient protrusion (white arrow) forming at the leading edge. Cell velocity fields obtained from particle image velocimetry of phase contrast images. (Scale bar, 100µm.) **C)** x-t kymograph of same region shown in B). **D-G)** Metrics from manual tracking of protrusions (circular assay format, compiled from 2 independent experiments). **D)** Number of protrusions per mm over 10 hours. Median values are 1.27 (n=7/5 monolayers for fast/slow, respectively). Two-tailed Mann Whitney test, p=0.0101. Error bars represent SD. **E)** Duration of protrusion tracks. (n=70/122 tracks for fast/slow, respectively). Two-tailed Mann Whitney test, p=0.1026. **F)** Track mean speed, microns per hour. (n=70/122 tracks for fast/slow, respectively). Two-tailed Mann Whitney test, p=0.0027. **G)** Persistence/confinement ratio of protrusion tracks. (n=70/122 tracks for fast/slow, respectively). Two-tailed Mann Whitney test, p=0.0883. **H)** Number of tracks over a 10-hour period plotted against relative area expansion, normalized to initial monolayer area.

### Competition between supracellular actomyosin cable and protrusion formation at the monolayer periphery dictates collective migration on viscoelastic substrates

To further investigate the dynamics of leader cell formation in our system, we looked more closely at the three-dimensional architecture of the leading edge. In line with prior studies, we observe a distinct multicellular actin cable along the periphery of monolayers on both fast- and slow-relaxing substrates, except in leader cells, which exhibit a lamellipodial protrusion at their leading edge. (23) (**Fig. 3 A and B, Movie S3)**. This cable is located at the basal plane of the monolayer, with rounded cells at the edge often appearing to spill out over the cable when viewed from above. (**Fig. 3B**). Monolayers are frequently thicker along the leading edge when a clear actin cable is present, indicating the possibility that the cable prevents forward movement and leads to cell pileup from the pressure of migration and/or proliferation in the rest of the monolayer (**Fig. 3C**). In contrast, monolayer heights within the bulk remain relatively constant, apart from local fluctuations at sites of cell division. In addition, supporting the view that the actin cable prevents forward movement and leads to cell-pile-up, is the observation that the cell edge becomes considerably flatter when a lamellipodial protrusion is present (**Fig. 3D)**.

**Figure 3.**
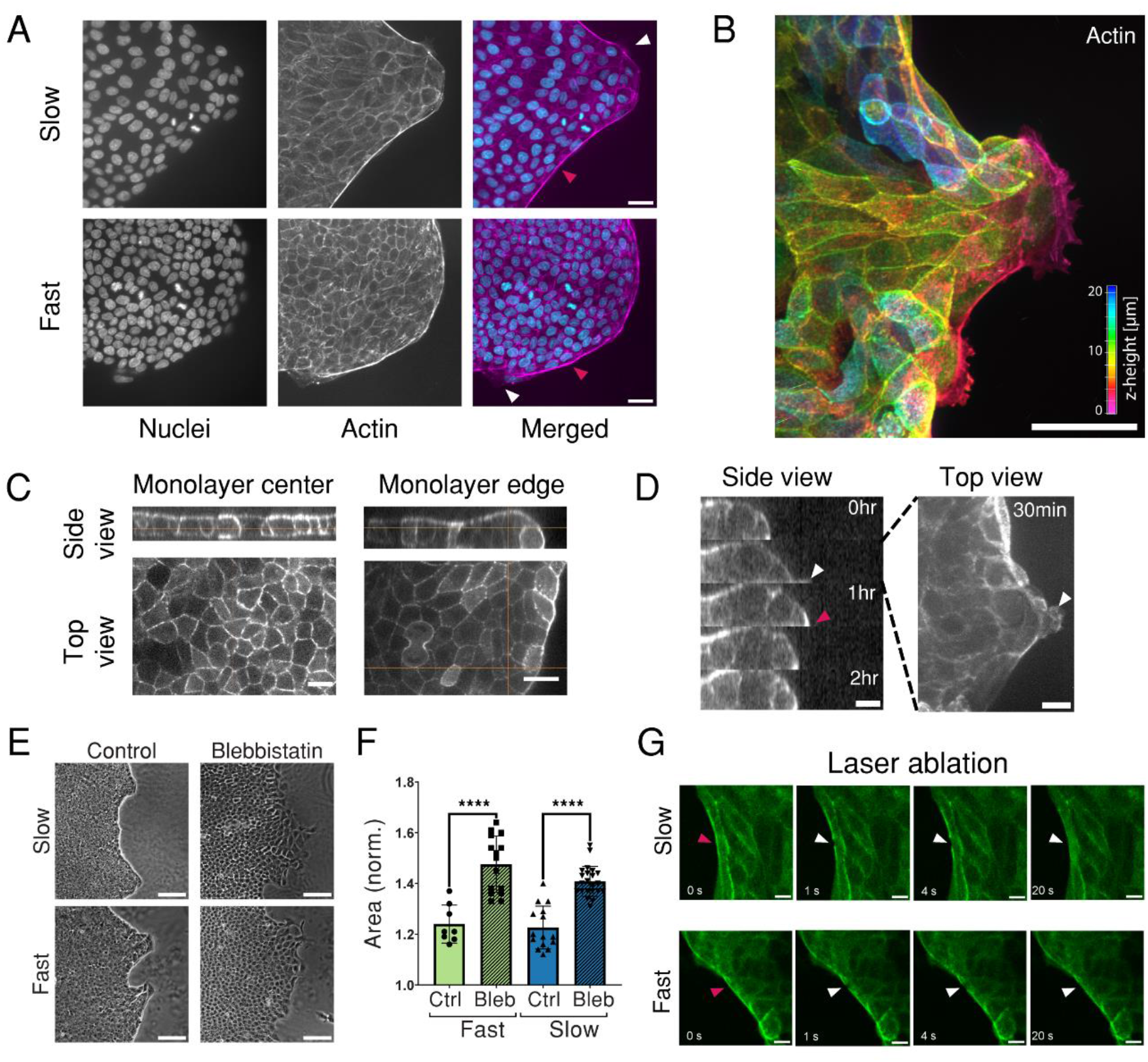
Competition between supracellular actomyosin cable and protrusion formation at the monolayer periphery. **A)** Monolayer leading edge with prominent actin cable (red arrows) interrupted by protrusions (white arrows). Hoechst and Phalloidin-488. (Scale bar, 25µm.) **B)** LifeAct-GFP volume rendering from z-stack, color-coded by z-depth (red is towards basal/substrate side, blue is towards apical side).(Scale bar, 50µm.) **C)** Monolayer height is uniform in the bulk and thicker at edges. LifeAct-GFP. (Scale bar, 10µm.) **D)** Monolayer edge temporarily flattens at location of protrusion (white arrow) and thickens when actin cable is re-established (red arrow). LifeAct-GFP. (Scale bar, 10µm.) **E)** Morphology of leading edge after addition of either DMSO (control) or blebbistatin. (Scale bar, 100µm.) **F)** Relative area expansion of radial clusters at 10 hours following addition of DMSO or blebbistatin. (n=8,15,15,22 monolayers, respectively, from two independent experiments) Tukey’s multiple comparisons test, p<0.0001. Error bars represent SD. **G)** Retraction of actin cable (white arrows) at site of laser ablation (red arrows). LifeAct-GFP. (Scale bar, 10µm.)

We next examined the dynamics of actin cable disruption and leader cell initiation. During transient bursts of protrusion, a peripheral cell will break past the actin cable to extend a prominent lamellipodia and drive a local forward surge in the monolayer. This nascent leader cell is often eventually pulled back in and the actin cable is re-established. (**Fig. 3D, Movie S4**) To globally disrupt the supracellular cable, as well as cell substrate contractility, we treated migrating monolayers with the myosin II inhibitor blebbistatin. (47–49) Blebbistatin addition led to significant roughening of the leading edge and acceleration of monolayer expansion on both fast- and slow-relaxing substrates. (**Fig. E and F, Movie S5**) Blebbistatin treatment is a blunt perturbation which may simultaneously impact several aspects of collective migration in addition to cable formation (e.g., cell-cell and cell-substrate force generation). Nevertheless, these results combined with examination of the leading-edge architecture suggest that restriction of protrusion formation by the supracellular actomyosin cable is a key feature of collective migration on viscoelastic substrates.

Prior studies have established that the supracellular actomyosin cable is under tension, which restricts new leader cell formation and enables the development of long fingers. (10) To probe cable tension in our system, we used laser ablation to cut a thin line across actin cables at various locations along the monolayer periphery. Following ablation, the severed cable edges quickly retract. The retraction speed and therefore cable tension appears to be variable depending on cell shape and proximity to leader cells, with faster retraction for straighter cables along elongated cells and closer to large multicellular protrusions. **(Fig. 3G, Fig. S3)**. The LifeAct signal recovers very quickly, appearing to reform the cable within tens of seconds. Interestingly, protrusion formation following laser ablation on short timescales (i.e., several minutes) is not observed, in line with recent experiments where the peripheral actin cable was locally disrupted. (50) It is possible that a more significant local perturbation of the cable could lead to leader cell formation on longer timescales in our system. Together, these data indicate that actin cables confine the monolayer, but that local severing of the actin cable is not sufficient to initiate a burst of protrusion.

### Cell motility in the monolayer bulk is enhanced on slow-relaxing vs fast-relaxing substrates

Given the similar responses thus far to perturbations of the actin cable on fast- and slow-relaxing substrates, we investigated additional features of collective migration which could more fully explain the differences in expansion between monolayers on the two substrates, and higher initiation of leader cells on the slower relaxing substrates. Collective migration is not solely driven by leader cells, but also involves active participation by cells within the monolayer bulk. While leader cells form easily visible lamellipodial protrusions at the monolayer periphery, the follower cells behind the leading edge have also been shown to extend cryptic lamellipodia underneath neighboring cells. (2, 51–54) Consistent with these previous data, cryptic lamellipodia and clear stress fibers in follower cells are observed at the basal plane of monolayers in our system. (**Fig. S4, Movie S6)**.

Focusing on the cells within the bulk, we observe that cell motility is enhanced in confluent regions on slow-relaxing substrates relative to fast-relaxing substrates. Monolayers on slow-relaxing substrates display more elongated cell shapes and dynamic movement, whereas monolayers on fast-relaxing substrates show characteristics of a jammed system, with fewer cell rearrangements and constrained motion. (**Fig. 4 A and B, Movie S7**) Accordingly, root mean square velocities within the monolayer bulk are higher on slow-relaxing substrates. (**Fig. 4C)**. In contrast, cellular organization within large multicellular protrusions appears fluid-like on both slow-relaxing and fast-relaxing substrates **(Fig. 4D)**.

**Figure 4.**
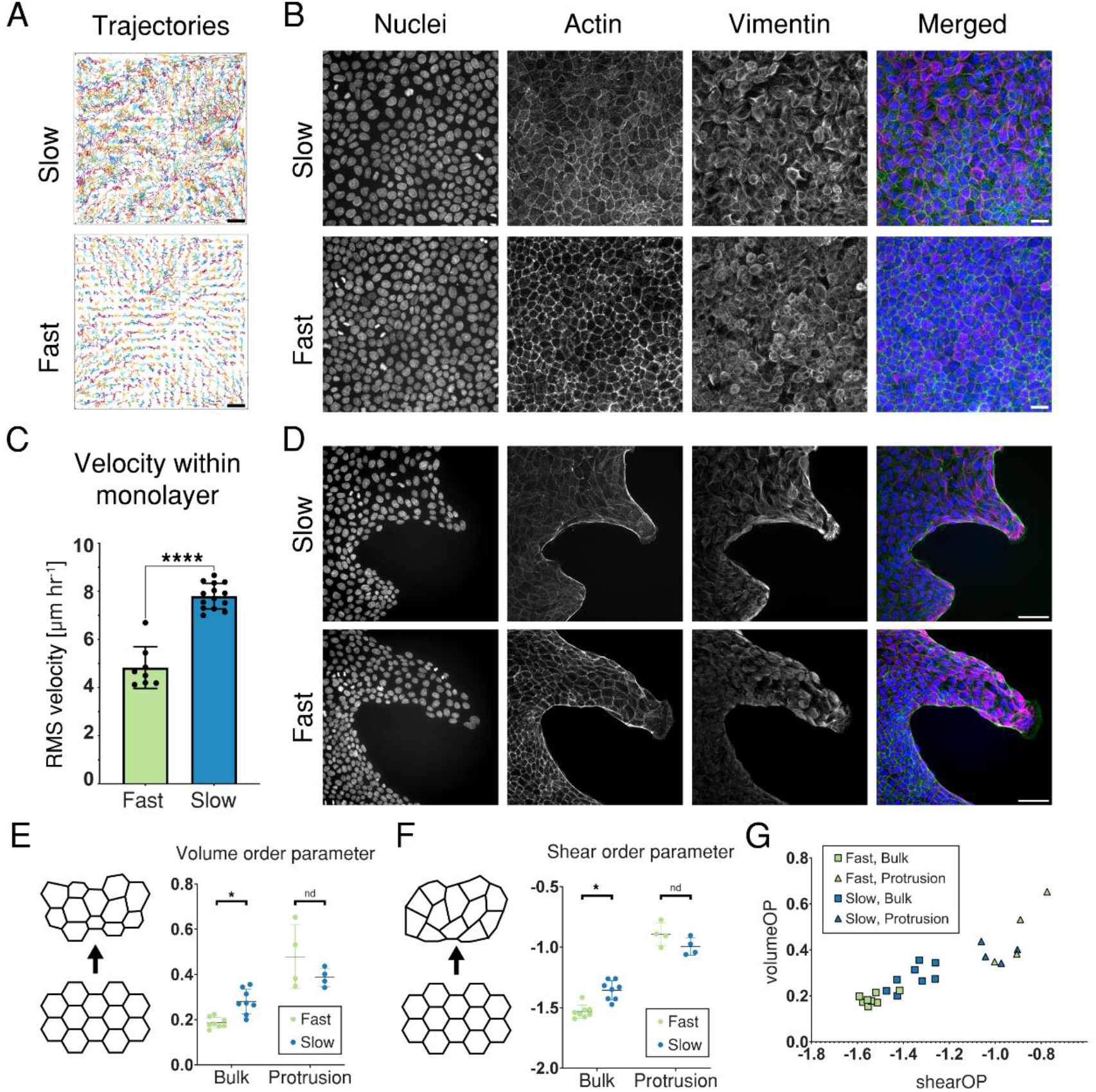
Cell motility in the monolayer bulk is enhanced on slow-relaxing vs fast-relaxing substrates. **A)** Representative cell trajectories over a 20-hour period, generated from cell velocity fields. (Scale bar, 25 µm.) **B)** Representative immunofluorescence images of regions within the monolayer bulk. (Scale bar, 20 µm.) **C)** RMS velocities within the bulk over a 20-hour period, measured for 250x250 µm regions taken from center of each monolayer, circular assay format. (n=8/14 clusters for fast/slow, respectively). **D)** Representative immunofluorescence images of well-developed protrusions along the monolayer edge. (Scale bar, 50 µm.) **E)** Volume order parameter (measure of variability in cell size) measured for images in the bulk or within multicellular protrusions. (n=8,8,4,4 images, respectively). Multiple Mann-Whitney tests, p=0.001 for bulk, p=0.486 for protrusions. **F)** Shear order parameter (value closer to 0 indicates higher shear of triangles) measured for the same set of images. Multiple Mann-Whitney tests, p=0.001 for bulk, p=0.200 for protrusions. **G)** Volume order parameter and shear order parameter plotted for all conditions. Error bars represent SD for all panels.

To characterize the spatial order of the system, we calculate volumetric and shear order parameters, (55) as well as an alignment parameter (q) to characterize anisotropy within the monolayer. (56) Cell areas and measures of variability in cell area (volume order parameter) and shear deformation (shear order parameter) are lower in the monolayer bulk on fast-relaxing substrates (**Fig. 4 E-G, Fig. S4**). On both substrates these parameters all increase significantly within protrusions. The shear order parameter in particular has previously been used to identify jamming transition points in confluent cell monolayers. (55) Together these data indicate that motility of cells within the monolayer bulk is enhanced on slow-relaxing substrates, whereas the confluent regions of monolayers on fast-relaxing substrates display characteristics of a jammed system.

### Substrate deformation by migrating epithelial monolayers is greater on faster-relaxing substrates

Finally, we sought to link monolayer motility more directly to viscoelastic properties of our alginate hydrogels by examining cell-substrate interactions. By tracking the position of fluorescent beads embedded within the alginate hydrogels, we observe significant deformation of the substrate directly in front of migrating leader cells, indicating that cells are physically coupled to the substrate and generating contractile forces (**Fig. 5A**). Substrate deformations are greater for protrusions on fast-relaxing substrates (**Fig. 5B**). We next observed that small groups of neighboring cells within the monolayer bulk transiently detached from the substrate following blebbistatin treatment on slow but not fast-relaxing substrates, suggesting that overall adhesion of monolayers to the substrate is weaker for slow-relaxing substrates (**Movie S7)**.

**Figure 5.**
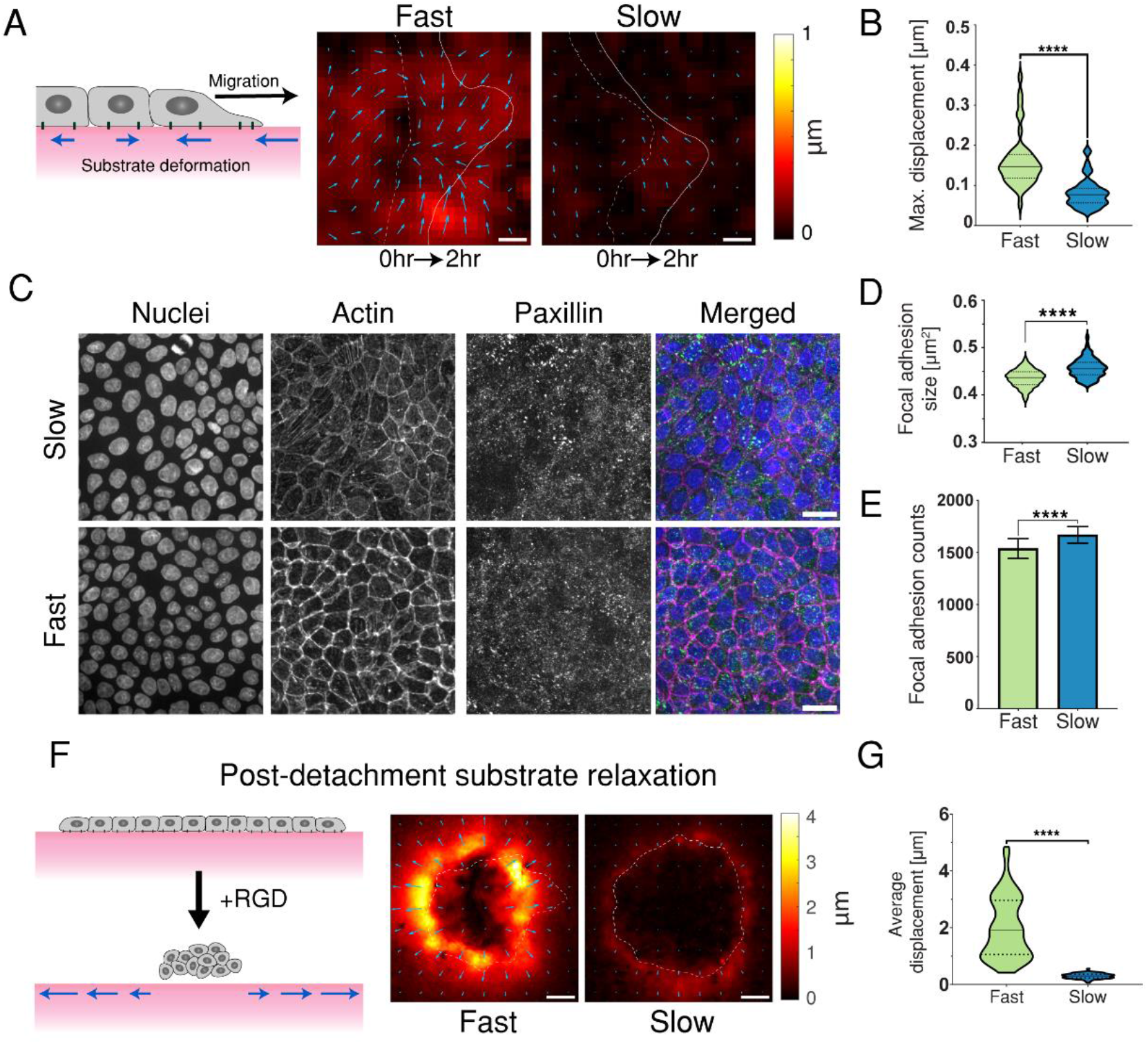
Substrate relaxation timescale alters deformation of viscoelastic substrates by epithelial monolayers. **A)** Substrate deformations generated by migrating protrusions over a two-hour interval. Initial and final positions of monolayer boundary indicated by dashed and solid lines, respectively. (Scale bar, 20 µm.) **B)** Maximum substrate deformation between successive frames, 20min interval. (n=43/46 protrusions for fast/slow, respectively, from two independent experiments). Two-tailed Mann-Whitney test, p<0.0001. **C)** Immunofluorescence images of paxillin adhesions for regions within the monolayer bulk. (Scale bar, 20 µm.) **D)** Size of paxillin foci. (n=67/77 for fast/slow, respectively, from two independent experiments). Two-tailed Mann-Whitney test, p<0.0001. **E)** Number of paxillin foci identified within a 330 by 330µm region in the monolayer bulk. (n=67/77 for fast/slow, respectively, from two independent experiments). Two-tailed Mann-Whitney test, p<0.0001. Error bars represent SD. **F)** Representative images of substrate relaxation immediately after monolayer detachment with RGD. Dashed line indicates monolayer boundary prior to detachment. (Scale bar, 200 µm.) **G)** Substrate relaxation after monolayer detachment with RGD. (n=55/79 monolayers for fast/slow, respectively, from five independent experiments. Two-tailed Mann-Whitney test, p<0.0001.

Prior single cell experiments and predictions from molecular clutch models demonstrated that cells on fast-relaxing substrates form bonds with greater lifetime and number. (34) The transient detachment of monolayers observed here could be explained by reduced formation of adhesions and/or faster timescales of adhesion turnover for slow-relaxing substrates. To directly visualize and quantify cell-substrate adhesions, we fixed monolayers after 24-48 hours of migration and imaged focal adhesion components, including vinculin and paxillin. Monolayers on both sets of hydrogels form focal adhesions throughout the bulk and within protrusions, without readily apparent differences in shape or distribution (**Fig. 5E, Fig. S5**). Quantification of paxillin foci within the monolayer bulk indicated only small differences in adhesion number and size between substrates (**Fig. 5 C and D)**.

To perturb cell-substrate adhesion more directly, we added soluble RGD peptide into the media after allowing monolayers to expand for 24 hours. Since RGD peptides coupled to the alginate polymer provide the only cell-adhesive ligand in this hydrogel system, addition of soluble RGD at sufficient concentrations would be expected to compete for integrin binding and interfere with cell adhesion. Following addition of 1mM RGD, portions of the monolayer edge begin to slowly retract, until a rapid cascade of retraction whereby monolayers curl up to a small fraction of their initial area and completely detach from the substrate (**Fig. S5**). This rapid retraction indicates that the entire monolayer is under tension, analogous to the response of a stretched elastic sheet pinned to a corkboard as pins are removed. Relaxation of the substrate immediately following monolayer release is significantly larger for fast-relaxing substrates, in line with results for protrusion-generated displacements (**Fig. 5 F and G**). Altogether, these results suggest that the timescale of substrate stress relaxation alters cell-substrate interactions to shape collective migration.

## Discussion

Here we found that substrate stress relaxation regulates collective cell dynamics on viscoelastic substrates. Epithelial clusters on slow-relaxing substrates show greater fluidity within the monolayer bulk than clusters on fast-relaxing substrates. Enhanced baseline motility on slow-relaxing substrates may provide the energy needed for cells at the periphery to overcome the barrier provided by the supracellular actin cable and extend more frequent lamellipodial protrusions. The emergence of protrusive leader cells in turn drives localized bursts of migration to accelerate overall monolayer expansion. In contrast, epithelial cells generate significant displacements on fast-relaxing substrates and adopt a more contractile, jammed state which slows down migration (**Fig. 6)**. This work adds to our understanding of tissue-substrate interactions and highlights the importance of dynamic material properties in collective cell migration.

**Figure 6.**
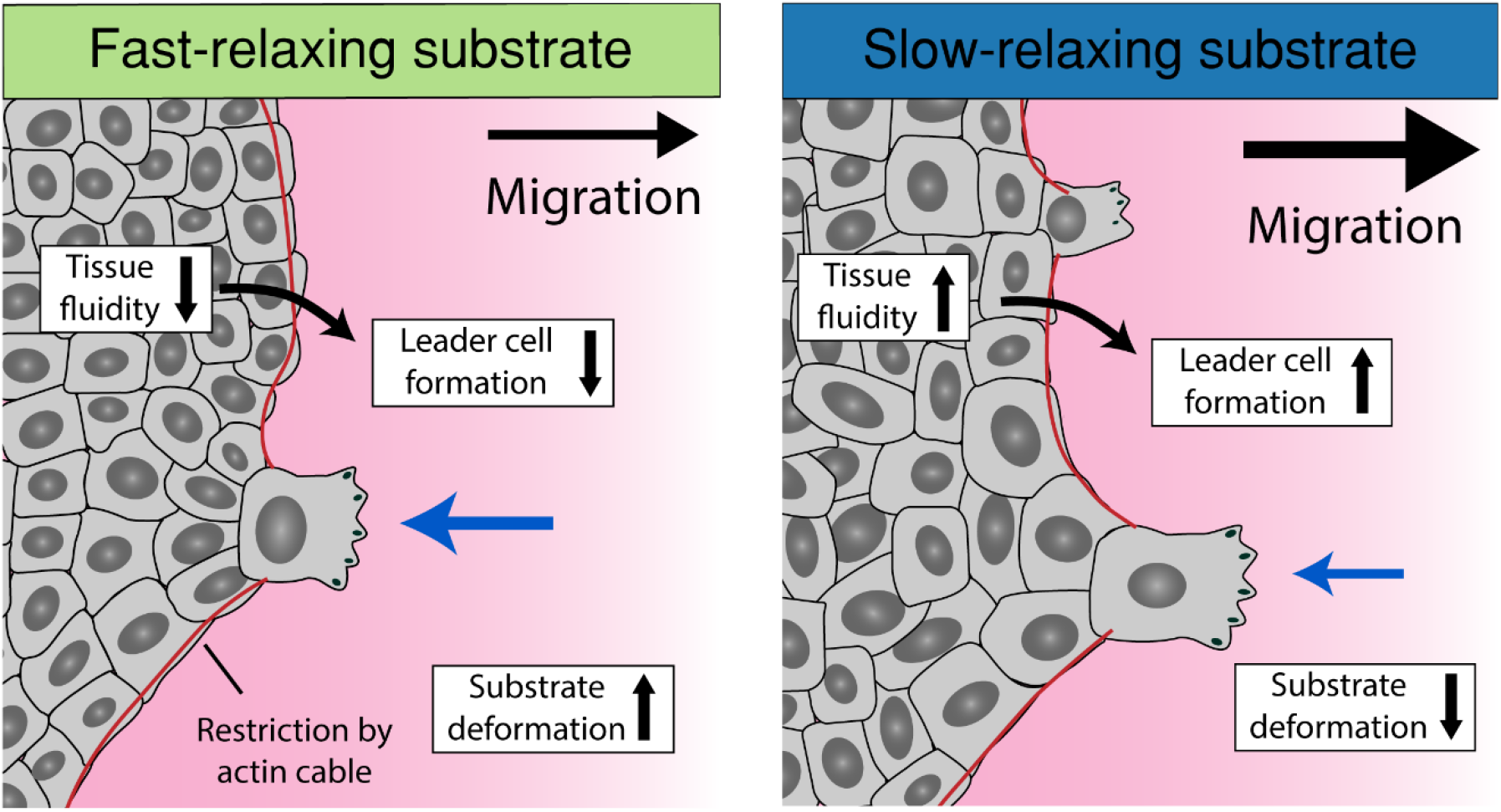
Monolayers on slow-relaxing substrates are more motile, deform the substrate less, and generate more frequent protrusive leader cells to collectively expand faster than monolayers on fast-relaxing substrates.

Despite increasing recognition that viscoelasticity is a key feature of many soft tissues, few studies have focused specifically on the role of substrate viscoelasticity in collective epithelial behavior. A recent study found that cancer cell clusters migrating on collagen substrates can generate local gradients in fiber density and stiffness to facilitate persistent directional motion, in a manner that is dependent on the timescale of substrate stress relaxation. (57) Additionally, investigations of epithelial monolayers on viscoelastic PDMS substrates have shown long-ranged correlation in collective motion and cell-type dependent coalescence of cell sheets, although the elastic and viscous components (storage and loss modulus) could not be independently tuned. (26, 58) One advantage of the alginate hydrogel system used in the present study is the ability to tune substrate stress relaxation timescales independently of substrate stiffness and cell-adhesive ligand density.

Here we observe faster expansion of MDCK monolayers on alginate substrates with equivalent stiffness and RGD ligand density, but with an order of magnitude difference in relaxation half-times. Our group recently reported the reverse trend between substrate relaxation timescale and single adherent cell migration in 2D, i.e., enhanced migration on faster-relaxing substrates. (34) Single cancerous and non-cancerous epithelial cells on soft, viscoelastic substrates migrated with rounded morphologies by extending thin filopodial protrusions, rather than the canonical lamellipodial protrusions observed on stiffer materials (e.g., glass). Relatedly, another study reported “viscotaxis” of mesenchymal stem cells (MSCs) cultured on polyacrylamide gels with a gradient of loss modulus, with the MSCs migrating towards the low loss modulus areas. (31) In contrast, collective migration of epithelial clusters in the present study is characterized by protrusive bursts of forward motion driven by leader cells with well-developed lamellipodia, in addition to a prominent supracellular cable restricting forward motion of the leading edge. This behavior bears resemblance to spontaneous polarization of single MDCK cells, in which cytoskeletal architecture at the cell edge depends on competition between two processes: Arp2/3-dependent growth of branched actin protrusions and myosin-dependent contraction of cables which suppresses polymerization by immobilizing actin within peripheral bundles. (49) Importantly, collective migration requires coordination of motion and force-generating processes across multiple cells. (59)

Lamellipodia-based cell crawling and purse-string like contraction of supracellular cables are two characteristic features of collective migration. Coordination of these migration modes in epithelial monolayers has been well-studied, particularly in the context of closure of small gaps. (15, 48, 60) Changes in actin flow and tension gradients due to physical or geometrical constraints enable large-scale curvature sensing to direct migration mode switching, with protrusions typically forming at positive (convex) curvature and cable formation at negative (concave) curvature. (17) Despite a global positive curvature initially opposed by the circular assay format, we observe distinct cable formation around the periphery of monolayers in line with similar studies on soft elastic substrates. (23) Cells at the edge of monolayers in our system periodically extend protrusions past this cable to initiate leader cell bursts with greater frequency on slow-relaxing substrates.

What collective phenomena might help to explain the dynamics of protrusion formation observed here? Prior work with small epithelial clusters breaking symmetry after release from confinement revealed that global cell alignment governed protrusion formation at weak points in an actomyosin cable distributed along the monolayer edge. (50) We have not specifically observed large scale alignment preceding leader cell formation in the present work, but we do find increased cell alignment within and close to established multicellular protrusions. Furthermore, we observe increased fluidity in the monolayer bulk on slow-relaxing substrates relative to fast-relaxing substrates. Unjamming in follower cells has been linked to greater fluctuations in cell-cell forces, which supports polarization and protrusion formation by leader cells. (47)

How then are emergent collective phenomena such as jamming influenced by substrate viscoelasticity? The molecular clutch model can provide a framework for understanding the relationship between motility, adhesion, and material relaxation timescales. (61, 62) A combination of modeling and experiments indicated that single cell spreading on soft substrates is maximized at an intermediate viscosity. (27) If the substrate relaxation timescale is faster than the timescale of clutch binding, the substrate is perceived as effectively softer, and spreading is decreased. If the substrate relaxation timescale is between the timescales of clutch binding and clutch lifetime, spreading is greatest because the bond experiences gradual relaxation of force after engagement. In the context of epithelial monolayers this could translate to longer cell-substrate adhesion lifetimes on substrates with faster stress relaxation, which could in turn produce a more jammed system with reduced fluidity. Indeed, a recent study of keratinocytes on nanopatterned substrates reported faster collective migration for conditions where focal adhesions were the most dynamic (i.e., shortest lifetimes). (63)

Adhesion dynamics may also impact protrusion formation at the leading edge more directly via adhesions between the substrate and the peripheral actin cable. Faster adhesion turnover on slow-relaxing substrates could result in a less homogenous cable structure which is more susceptible to interruption by lamellipodial protrusions. (60) Using substrates with tunable RGD density, prior work established a link between reduced cell-substrate adhesion, weakened actin belts, and a higher frequency of lamellipodia formation, a combination of features which resembles the slow-relaxing substrate condition in the present study. (64) Alternatively, stronger adhesion between the cable and the substrate on fast-relaxing substrates could impede forward movement of the leading edge and increase cell crowding within the monolayer.

Importantly, viscoelastic substrates exhibit creep under static loading, and viscosity of the basal lamina underlying *in-vivo* epithelia on timescales relevant to collective migration has been suggested to play a role in long-ranged correlated motion of cell sheets. (26) During closure of small gaps in a confluent epithelium, contraction of a supracellular actomyosin ring can generate substrate displacements directed towards the damaged area to accelerate wound healing, with greater impact for softer substrates. (15) In contrast, substrate displacements generated by expanding monolayers point in the opposite direction of migration and may instead work against expansion, particularly for soft, viscoelastic materials. Substrate flow may therefore play an important and context-dependent role in shaping tissue dynamics.

Ultimately, the effect of substrate stress relaxation on collective epithelial behavior is likely dependent on the balance between cell-cell and cell-substrate forces resulting from particular combinations of intrinsic cell properties, including baseline contractility and cell-cell adhesion strength, and extrinsic features of the environment, such as substrate stiffness and relaxation timescales. Our understanding of these relationships will continue to benefit from careful consideration across multiple scales, from molecular mechanotransduction and adhesion dynamics to the coordination of motion and forces across distances far greater than the size of a single cell.

## Materials and Methods

### Cell culture

Parental MDCK type II cells (gift from Dr. William J. Nelson, Stanford University) were grown in low-glucose DMEM (11885084, Gibco) supplemented with 10% FBS (SH30071.03; GE Healthcare) and 100 U/ml penicillin-streptomycin (15140; Thermo Fisher Scientific). Growth media for MDCK cells stably expressing LifeAct-GFP (gift from Dr. Jens Möller, ETH Zurich, Zurich, Switzerland) were additionally supplemented with 250 µg/ml geneticin (10131027; Thermo Fisher Scientific).

### Alginate hydrogel preparation

Sodium alginates were purchased from FMC BioPolymer (LF20/40; 280 kDa MW) and Dupont Corporation (ProNova UP VLVG; 28 kDa MW) and prepared as described previously. (38) RGD peptides were coupled to alginate using carbodiimide chemistry to reach a final peptide concentration in the alginate gels of 1500μm, as described previously. (35, 36, 65) Lyophilized alginate was reconstituted at 3% w/v in DMEM (11885084, Gibco). Alginate was mixed with a solution of DMEM and calcium sulfate to form ionically crosslinked hydrogels. The concentration of calcium sulfate was adjusted in order to vary the initial elastic modulus of the hydrogels.

To prepare thin alginate gels for timelapse imaging of epithelial monolayers, spacer rings with 19mm outer diameter and 14mm inner diameter were cut from sheets of .010”(∼250μm-thick) silicone (Specialty Manufacturing, Inc) and placed in glass-bottom dishes (D35-20-1.5H, Cellvis). Alginate solutions were mixed and quickly deposited into glass dishes before gently pressing round 18-mm coverslips down onto the ring spacers to generate 250μm-thick hydrogels. To improve adhesion of the thin alginate gels to the dish and prevent drift during imaging, the glass microwells were incubated with 0.01x poly-l-lysine (PLL) for 30 minutes before hydrogel preparation. The alginate gels were covered with media and left overnight in the cell culture incubator to equilibrate.

### Alginate hydrogel mechanical characterization

Unconfined compression tests were performed on a 5848 MicroTester (Instron) to characterize the initial elastic modulus and stress relaxation halftime for alginate hydrogel disks with a diameter of 8mm and a thickness of 2mm. (30) Briefly, disks were compressed to 15% strain at a deformation rate of 1mm/min and held at 15% strain until the end of the test. The initial elastic modulus was measured by taking the slope of the stress-strain curve between 5% and 10% strain, and stress relaxation halftime was measured as the time required for the peak stress at 15% hold strain to drop to half of its initial value. Tests were performed on at least 3 biological replicates per alginate formulation.

### Micropatterning epithelial monolayers

Two different assay formats were used for stencil-based micropatterning of epithelial monolayers on thin alginate gels. For the rectangular assay format, magnetic PDMS stencils were cast from 3D-printed molds (Protolabs) with 500μm-wide rectangular features. Magnetic PDMS was created by mixing equal parts (by weight) of magnetic powder and PDMS (Kraden; Sylgard 184). For the circular assay format, PDMS stencils were cast from 3D-printed molds with 700μm-diameter circular features. In this format, a ring-shaped gasket (19mm outer diameter, 12mm inner diameter) made of magnetic PDMS was used to hold the non-magnet stencil in place during the course of cell seeding and incubation before stencil removal. All stencils were passivated before used by overnight incubation at 37°C in 10% bovine serum albumin (A4503; Sigma). Stencils were rinsed 2x in PBS and dried before placing on gels for cell seeding. A small volume of cell suspension was added to each gel to achieve a seeding density of ∼300*10^3^ cells per cm^2^. Dishes were placed on a custom magnet holder and the entire assembly was placed in the incubator overnight. After monolayers reached confluence, stencils were carefully removed with tweezers and the gel surface was gently rinsed with cell culture media to remove stray cells before beginning experiments. For inhibitor experiments, monolayers were released from confinement and allowed to migrate for 6hrs before replacing with media containing either 50μM blebbistatin or DMSO (control).

### Live timelapse microscopy of epithelial monolayers

Timelapse microscopy was performed on a Nikon ECLIPSE Ti2 inverted microscope with a Crest XLight spinning disk confocal unit, with 10x, 10x phase, 20x and 40x objectives (numerical aperture/NA of 0.45, 0.3, 0.75, and 1.15 respectively). Cells were maintained at 37°C and 5% CO2 for all live-cell experiments. To image monolayers with higher magnification (i.e., using the 40x objective), some alginate gels were prepared on free-standing coverslips and then flipped over onto a spacer in a glass bottom dish, such that the fluorescent illumination light path passed through one layer of glass and then a small volume of culture media, as opposed to passing through 250μm of alginate before reaching the cells.

### Laser ablation

Laser ablation was performed on a laser scanning confocal microscope (Zeiss LSM 780) at the Stanford Cell Sciences Imaging Facility using a Mai Tai DeepSee (Spectra Physics) multiphoton laser (800-nm wavelength), 20×PLAN APO NA 10.8 air immersion objective and Zen Black software. MDCK cells stably expressing LifeAct-GFP were maintained in cell culture conditions at 37°C with 5% CO2. Ablation consisted of a single line scan 0.1μm wide with a pixel dwell time of 108μs.

### Immunofluorescence imaging of fixed monolayers

Monolayers were fixed with 4% paraformaldehyde for 10 min and then washed 3 times with tris-buffered saline (TBS) containing 10 mM CaCl2 (cTBS) to preserve alginate hydrogel integrity. 150 µL of 1% agarose (Sigma, A9045) was deposited on top of the hydrogel to prevent monolayers from detaching. Fixed samples were permeabilized with 0.5% Triton X-100 (Sigma) in cTBS for 30 minutes and blocked with 1% BSA, 10% goat serum, 22.52 mg/mL glycine in cTBST (cTBS+ 0.1% Tween 20) for 1 hour. Primary antibodies targeting paxillin (Y113, Abcam ab32084) were used with together with Alexa-488 conjugated secondary antibody targeting rabbit (Invitrogen, A11008). Nuclei and actin were visualized with Hoechst 33342 (Invitrogen, H3570) and Phalloidin-555 (Invitrogen, A34055), respectively. All antibodies were diluted in cTBST and incubated for 3 hours in 4C. Samples were washed overnight in cTBST after addition of primary antibodies and for 3 hours before image acquisition. Images were acquired using a Nikon CFI Apo LWD 40X/1.15 NA water immersion objective.

### Image analysis

To obtain monolayer areas to quantify expansion and edge velocities, monolayer boundaries were segmented either using a custom Matlab script or a semi-automated thresholding method in ImageJ. For analysis of cell velocity fields and substrate displacements, a custom Matlab image analysis pipeline was adapted from scripts by Jacob Notbohm (University of Wisconsin-Madison).

(45) These scripts make use of a fast iterative digital image correlation (FIDIC) algorithm created by the lab of Christian Franck (University of Wisconsin – Madison). (66) For fluidity quantification, Delaunay triangulation of segmented cell nuclei from fixed monolayers was used to calculate shear and volume order parameters, which characterize the average degree of amorphousness and density variation in the tissues, respectively. (55, 67, 68) For adhesion quantification, images were adjusted to have the same brightness by histogram matching and the Focal Adhesion Analysis Server (FAAS) was used with a threshold of 2. (69)

## Acknowledgments

F.C. acknowledges support from a National Science Foundation Graduate Research Fellowship (DGE-1656518). O.C. acknowledges support from a National Science Foundation CAREER award (CMMI 1846367) and a National Institutes of Health National Cancer Institute grant (R37 CA214136). We also thank members of the Chaudhuri lab, especially Aashrith Saraswathibhatla and Dhiraj Indana, for their assistance and advice.

