## Supplemental Appendix for "Substrate stress relaxation regulates monolayer fluidity and leader cell formation for collectively migrating epithelia"

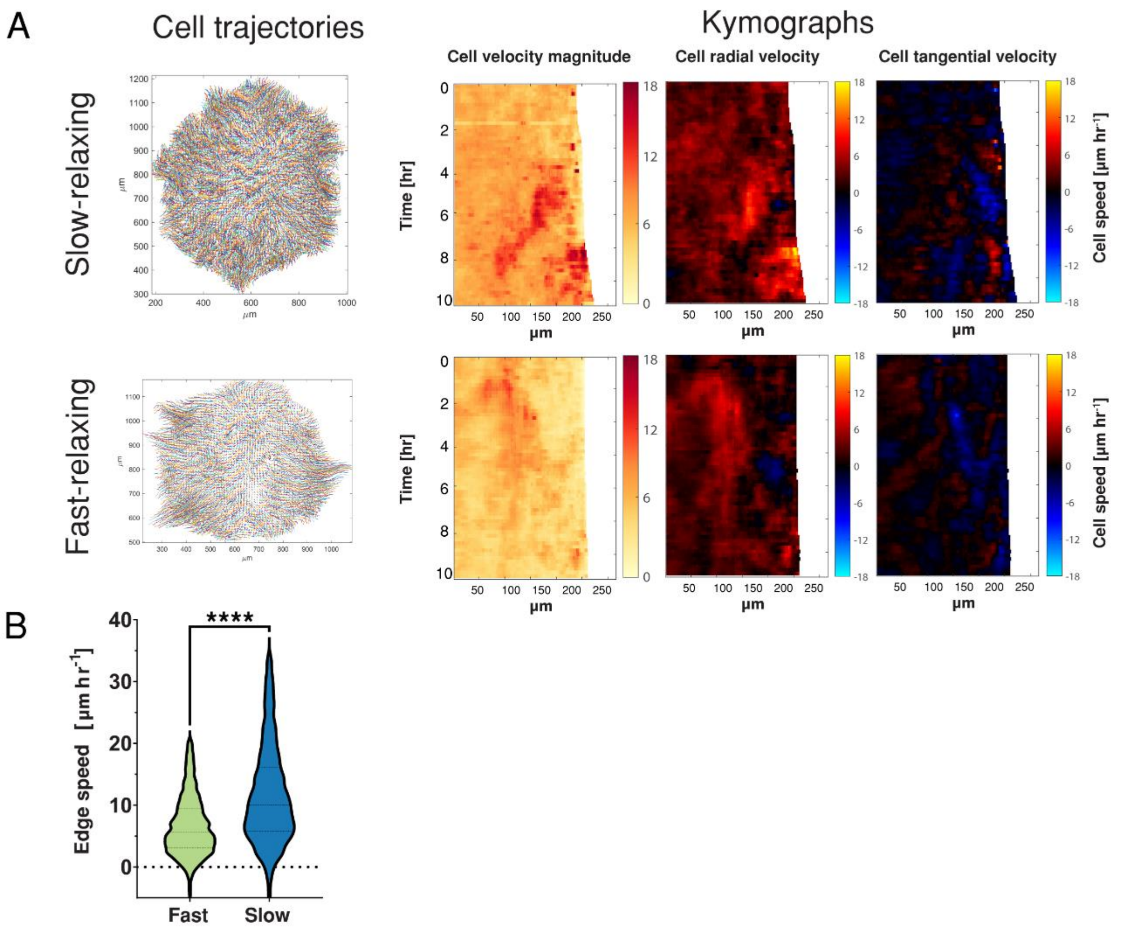


**Fig. S1. Epithelial monolayers expand faster on slow-relaxing vs fast-relaxing substrates.** **A)** Cell trajectories and kymographs for radial clusters over 10 hours of migration, from particle image velocimetry (see methods). **B)** Edge velocities for rectangular assay format. (n=6397/3541 tracks from 12 monolayers each for fast/slow, respectively, from two independent experiments). Two-tailed Mann-Whitney test, $p<0.0001$.

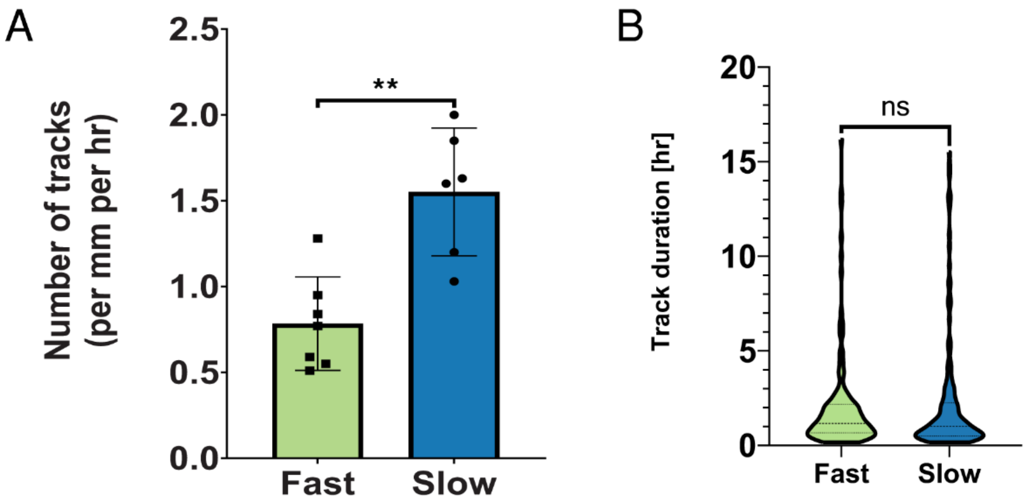


**Fig. S2. Protrusion quantification for rectangular clusters. A)** Number of tracks per mm per hr. (n=7/6 monolayers for fast/slow-relaxing, respectively). Two-tailed Mann-Whitney test, p=0.0047. **B)** Track duration. (n=150/157 tracks for fast/ slow, respectively). Two-tailed Mann-Whitney test, p=0.3319.

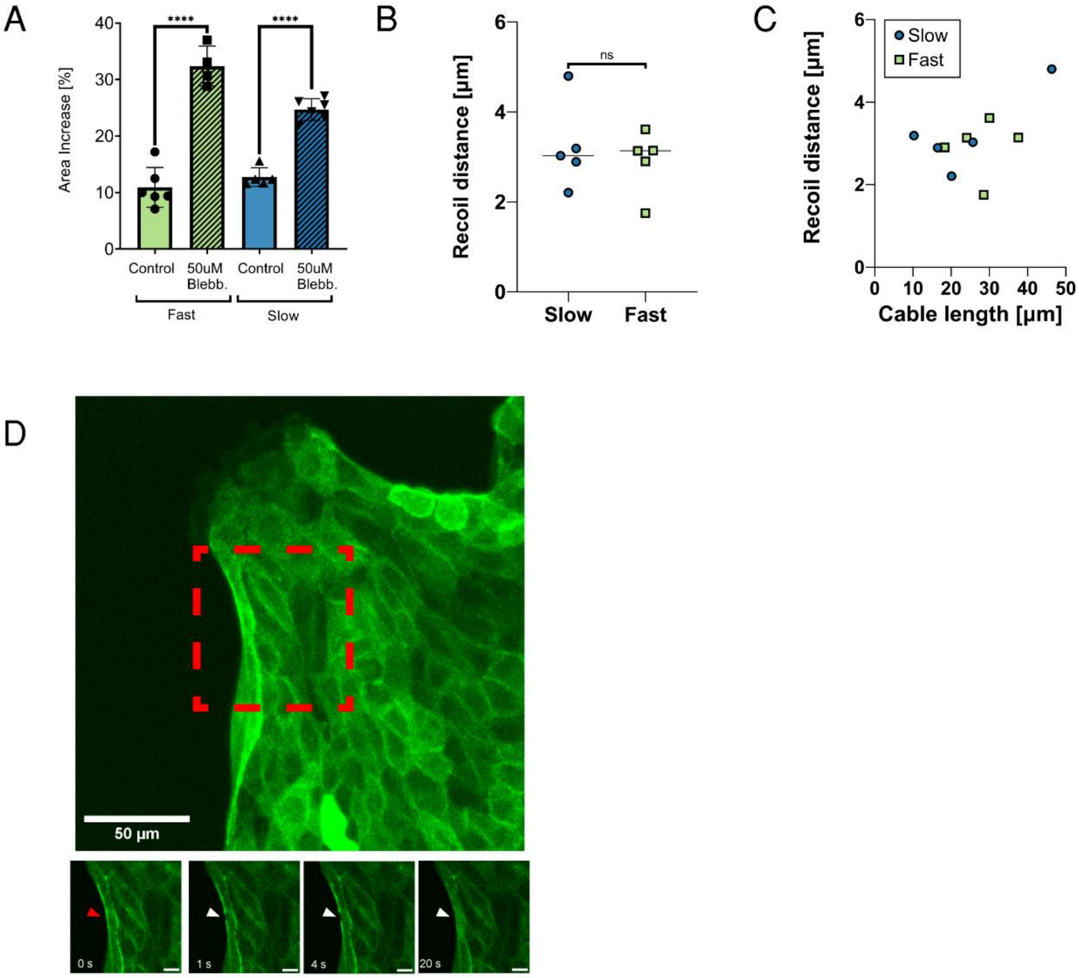


**Fig. S3. A)** Relative area expansion of rectangular clusters at 6 hours following addition of DMSO or blebbistatin. (n=6,4,5,6 monolayers, respectively). Sidak's multiple comparisons test, $p<0.0001$. **B)** Recoil distance of actin cable following laser ablation, measured 1s post-ablation. (n=5/5 for slow/fast, respectively). Two-tailed t-test, $p=0.57$. **C)** Recoil distance plotted against the initial length of the actin filament along the cell edge. **D)** Leader cell close to an ablated actin cable, for one of the ablations shown in Figure 3. Retraction of actin cable (white arrows) at site of laser ablation (red arrows). LifeAct-GFP. (Scale bars, 50µm for top panel, 10µm for bottom panels).

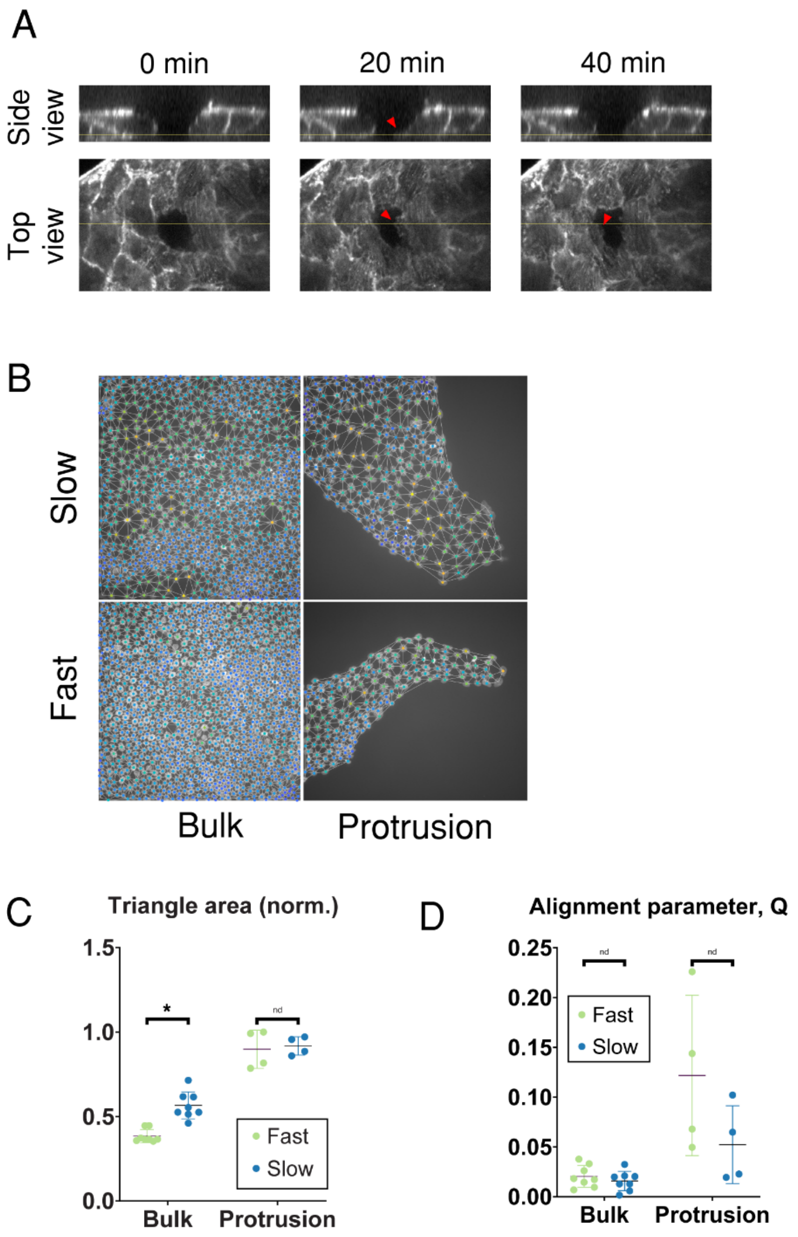


**Fig. S4. A)** Cryptic lamellipodia (red arrows) visible extending underneath cell lacking LifeAct-GFP expression. **B)** Triangulation of segmented nuclei used to characterize cell density, shear, and alignment. **C)** Triangle area (analogous to cell size), measured for images in the bulk or within multicellular protrusions. (n=8,8,4,4 images, respectively). Multiple Mann-Whitney tests, p=0.0002 for bulk, p>0.999 for protrusions. **D)** Alignment parameter (Q) measured for the same set of images. Multiple Mann-Whitney tests, p=0.505 for bulk, p=0.200 for protrusions.

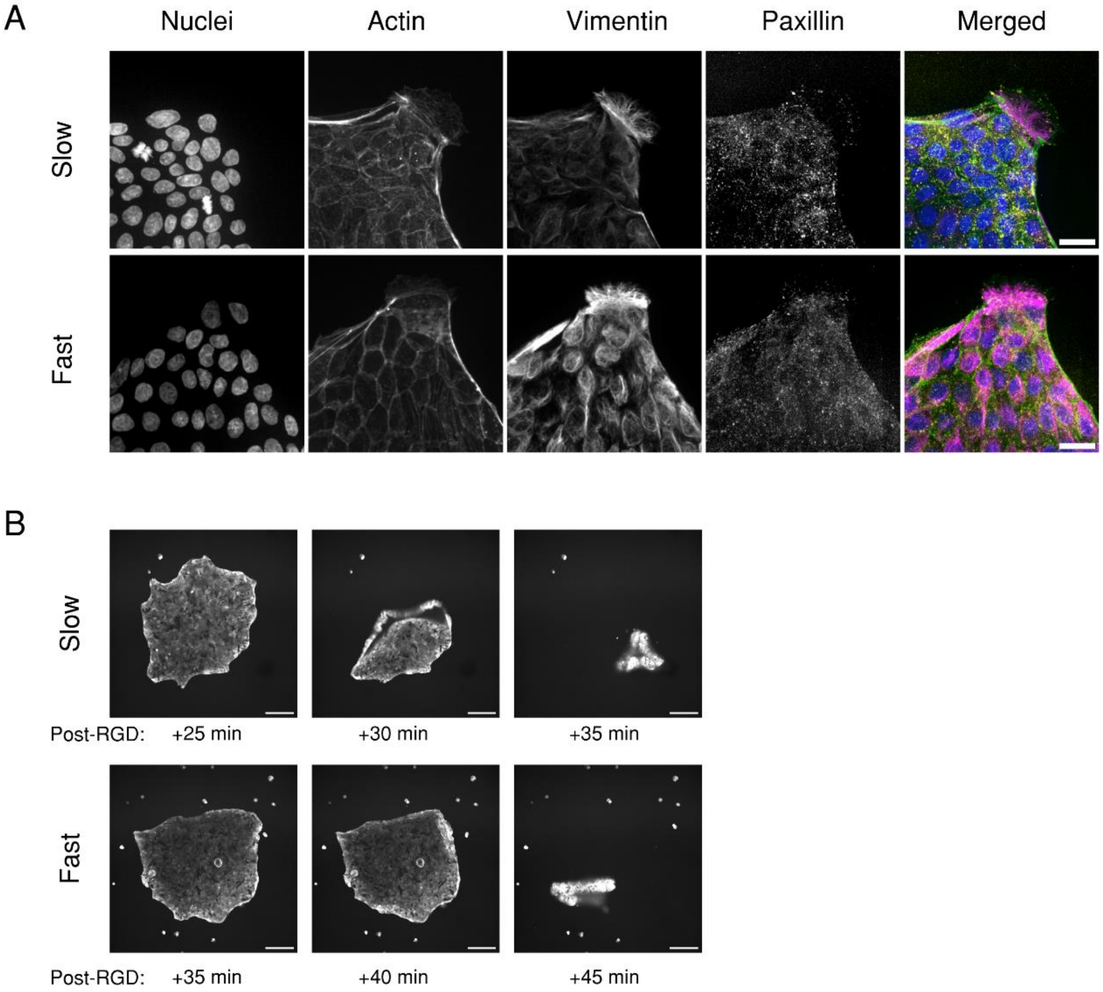


**Fig. S5. A)** Immunofluorescence images of adhesions underneath protrusions along the monolayer edge. (Scalebar, 20 μm). **B)** Time sequence of monolayer detachment following addition of 1mM RGD. LifeAct-GFP. (Scalebar, 200 μm).

**Movie S1.** Transient bursts of migration at leading edge of monolayer. LifeAct GFP. 10 min/frame. Scale bar = 100 µm.

**Movie S2.** Protrusion tracking on slow-relaxing substrate. Lines represent tracks, color-coded by protrusion duration. 10 min/frame. Scale bar = 100 µm.

**Movie S3.** 3D rendering of the leading edge of a monolayer, with prominent leader cell and lamellipodia. LifeAct GFP. Color coded by z-depth: blue at apical surface to red at basal surface.

**Movie S4.** 3D rendering of transient protrusion formation. LifeAct GFP. Color coded by z-depth: green at apical surface to red at basal surface. 10 min/frame.

**Movie S5.** Leading edge of monolayers with DMSO (control) or blebbistatin added at 5hrs, on fast- and slow-relaxing substrates. Scale bar = 100 µm.

**Movie S6.** Cryptic lamellipodia on fast-relaxing substrates. LifeAct GFP at apical and basal planes of monolayer. 10min/frame. Scale bar = 20 µm.

**Movie S7.** Bulk with control (DMSO) or blebbistatin added at 5hrs, on fast- and slow-relaxing substrates. 10min/frame. Scale bar = 100 µm.
