## Supplementary figures and images for "Substrate stress relaxation regulates monolayer fluidity and leader cell formation for collectively migrating epithelia"

### Movie S1

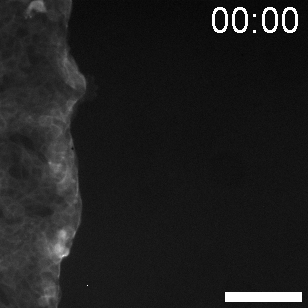

### Movie S2

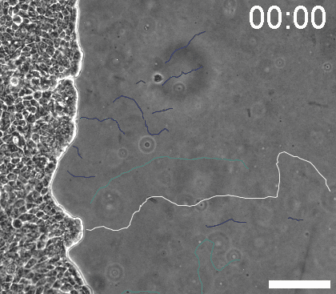

### Movie S3

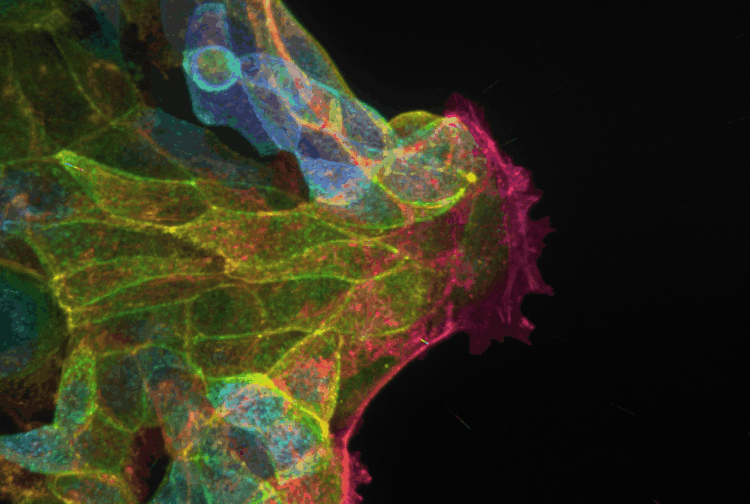

### Movie S4

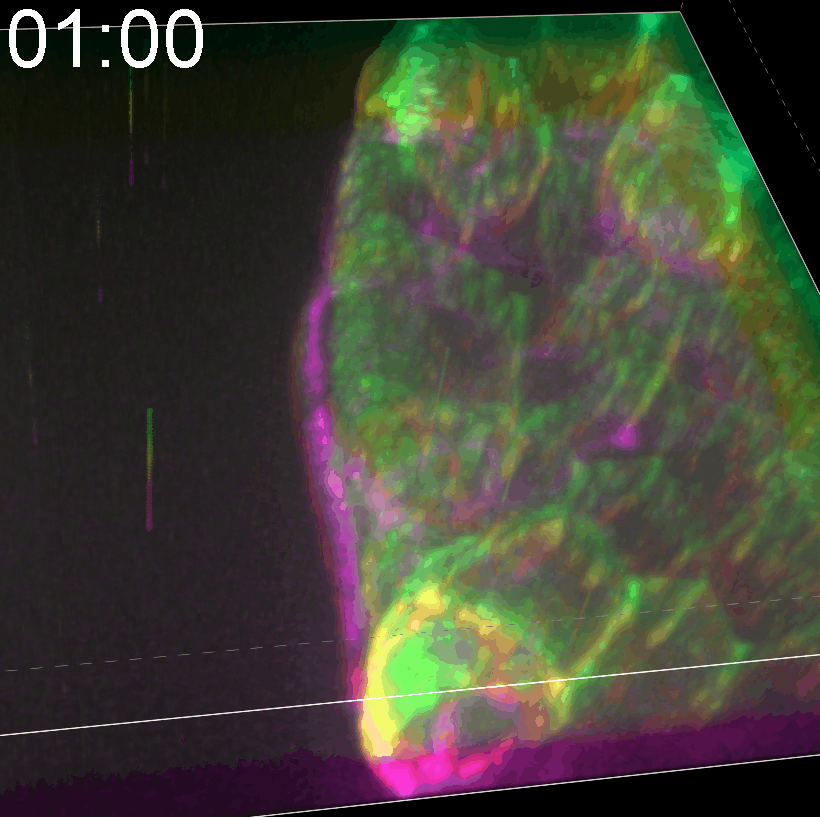

### Movie S6

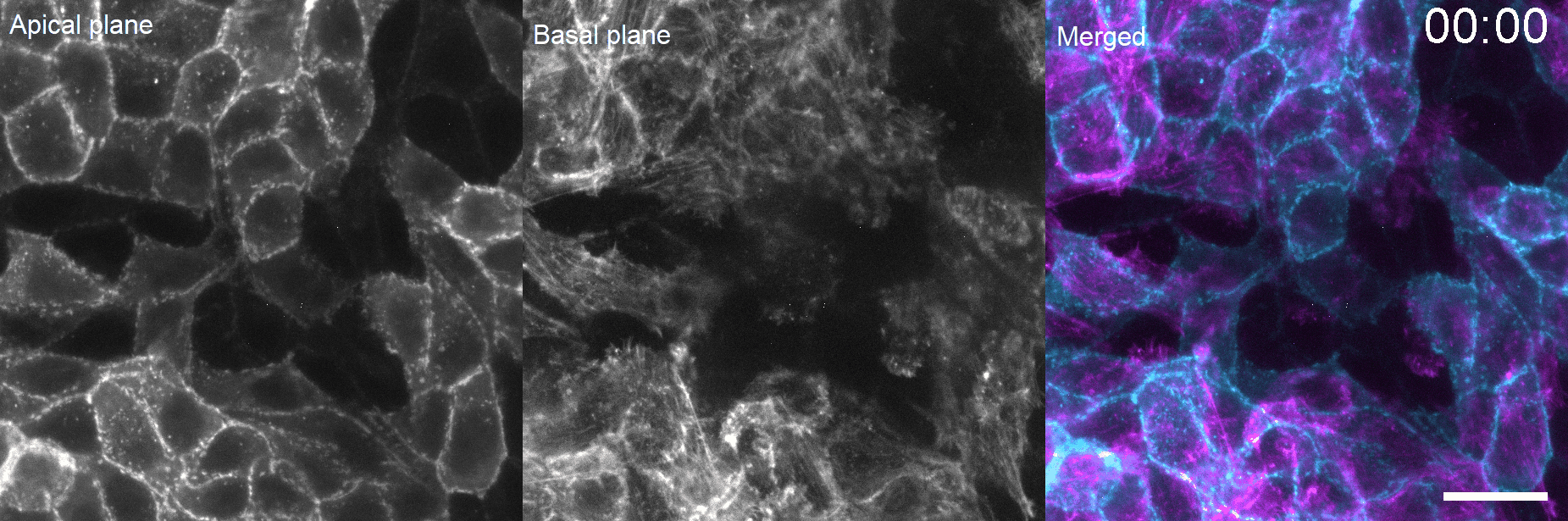
